# HTS-Oracle v2: Prospective AI-Guided Discovery and Experimental Validation of Small Molecule Modulators Across Multiple Targets

**DOI:** 10.64898/2026.06.15.732399

**Authors:** Somaya A. Abdel-Rahman, Moustafa T. Gabr

## Abstract

High-throughput screening (HTS) remains the cornerstone of early-phase small molecule discovery yet consistently underperforms against immunotherapy targets, yielding validated hit rates below 0.1%. Here we introduce HTS-Oracle v2, which features rigorous cross-validation that ensures honest performance estimates. HTS-Oracle v2 was trained and validated across four clinically significant immune checkpoint targets (CD28, ICOS, LAG-3, and TIGIT) achieving ROC-AUC values of 0.968, 0.969, 0.875, 0.928 respectively under rigorous cross-validation. For prospective experimental validation, HTS-Oracle v2 was applied to an 8,960-compound Enamine Protein Mimetic Library, selecting only 25 compounds per target for experimental testing using temperature-related intensity change (TRIC) technology, a 99.7% reduction in screening burden. HTS-Oracle v2 identified 4, 5, 4, and 6 validated binders from 25 prospectively selected compounds per target, corresponding to validated hit rates of 16%, 20%, 16%, and 24%, respectively. Notably, 67-80% of all experimentally confirmed hits across the full 8,960-compound library were captured within just 25 model-selected compounds per target. For CD28, this represents a 28-fold improvement over HTS-Oracle v1 (239× versus 8.4×), establishing HTS-Oracle v2 as an efficient platform for AI-guided prospective hit discovery across immunotherapy targets.

## Introduction

High-throughput screening (HTS) remains the cornerstone of early-phase small molecule drug discovery, yet consistently underperforms in practice, yielding hit rates of 1-2% under favorable conditions and frequently below 0.3% for challenging targets, while imposing enormous resource burdens in protein, consumables, and instrumentation time.^1,2^ These inefficiencies are especially pronounced against immune checkpoint targets, which include both inhibitory receptors such as lymphocyte activation gene-3 (LAG-3) and T cell immunoreceptor with immunoglobulin and ITIM domains (TIGIT), and costimulatory receptors such as cluster of differentiation 28 (CD28) and inducible T cell costimulator (ICOS).^3,4^ These proteins play central roles in T cell activation and antitumor immunity, and their therapeutic validation has been transformative, the FDA approval of relatlimab/nivolumab for LAG-3/PD-1 co-blockade in advanced melanoma,^5^ alongside active clinical programs targeting TIGIT,^6^ ICOS,^7^ and CD28,^8^ underscores the opportunity at this target class. However, immune checkpoints predominantly operate through protein-protein interaction (PPI) interfaces characterized by large, shallow, and featureless binding surfaces that present remarkable challenges to conventional small molecule discovery,^9,10^ contributing to the extremely low hit rates observed experimentally.^11^

Artificial intelligence (AI) and machine learning have emerged as powerful strategies to address these limitations, enabling large-scale virtual screening and compound prioritization from vast chemical spaces.^12,13^ Ensemble architectures combining transformer-based molecular language models such as ChemBERTa^14^ with traditional cheminformatics descriptors including Morgan fingerprints, MACCS keys, and physicochemical properties have shown strong performance across diverse bioactivity prediction tasks.^15,16^ However, a persistent limitation of many published AI-guided virtual screening platforms is reliance on random cross-validation splits during model development.^17,18^

Our group has established a sustained research program targeting immune checkpoints with small molecules, reporting first-in-class inhibitors of multiple checkpoints identified through complementary approaches including pharmacophore-based virtual screening, affinity selection-mass spectrometry, and TRIC-based high-throughput screening.^19–26^ These efforts have validated the experimental infrastructure and biological rationale that underpin HTS-Oracle, a retrainable multimodal deep learning platform that achieved up to 176-fold enrichment and greater than 99% reduction in screening burden for TREM2 and CHI3L1.^27^ However, HTS-Oracle v1 achieved 8.4-fold enrichment for CD28, its most challenging target, reflecting the severe class imbalance and structural complexity inherent to CD28’s protein–protein interaction interface.^27^

Several studies have demonstrated the value of computational hit discovery approaches through prospective experimental validation of model-selected compounds, including deep learning-guided discovery campaigns and large-scale virtual screening efforts.^28–32^ Despite these advances, relatively few studies have prospectively evaluated AI-guided hit discovery across multiple immune checkpoint targets within a unified experimental framework. Here, we present HTS-Oracle v2, the next generation of our AI-guided hit discovery platform, and evaluate its prospective performance across four therapeutically relevant immune checkpoint targets (CD28, ICOS, LAG-3, and TIGIT). Building upon HTS-Oracle v1, which demonstrated the feasibility of AI-guided hit enrichment for individual targets, HTS-Oracle v2 expands prospective deployment to a broader panel of immune checkpoint proteins. To assess its real-world utility, we prospectively screened an 8,960-compound Enamine Protein Mimetic Library and experimentally validated model-selected compounds through orthogonal binding and functional assays. HTS-Oracle v2 recovered 67-80% of all experimentally validated library hits while achieving enrichment factors of 239–286-fold over full-library random screening, including a 28-fold improvement in enrichment for CD28 relative to HTS-Oracle v1, demonstrating the utility of AI-guided prioritization for accelerating the discovery of functional small-molecule immune checkpoint modulators.

## Results and Discussion

### Dataset Curation and Rigorous Validation Design

Training datasets were compiled from ChEMBL and HTS studies for four immune checkpoint targets: CD28, ICOS, LAG-3, and TIGIT. The curated libraries comprised 10,691 compounds for CD28, 10,684 for ICOS, 4,610 for LAG-3, and 4,937 for TIGIT (Figure 1A). Scaffold diversity, measured by unique Murcko scaffolds, was 3,833, 3,832, 2,230, and 2,186 respectively, confirming broad chemical space coverage across all four targets. The two larger datasets, CD28 and ICOS at approximately 10,700 compounds each, reflect extensive ChEMBL curation supplemented by published screening data, while LAG-3 and TIGIT represent smaller but pharmacologically rich collections consistent with the more recently established therapeutic relevance of these emerging checkpoints.

**Figure 1.**
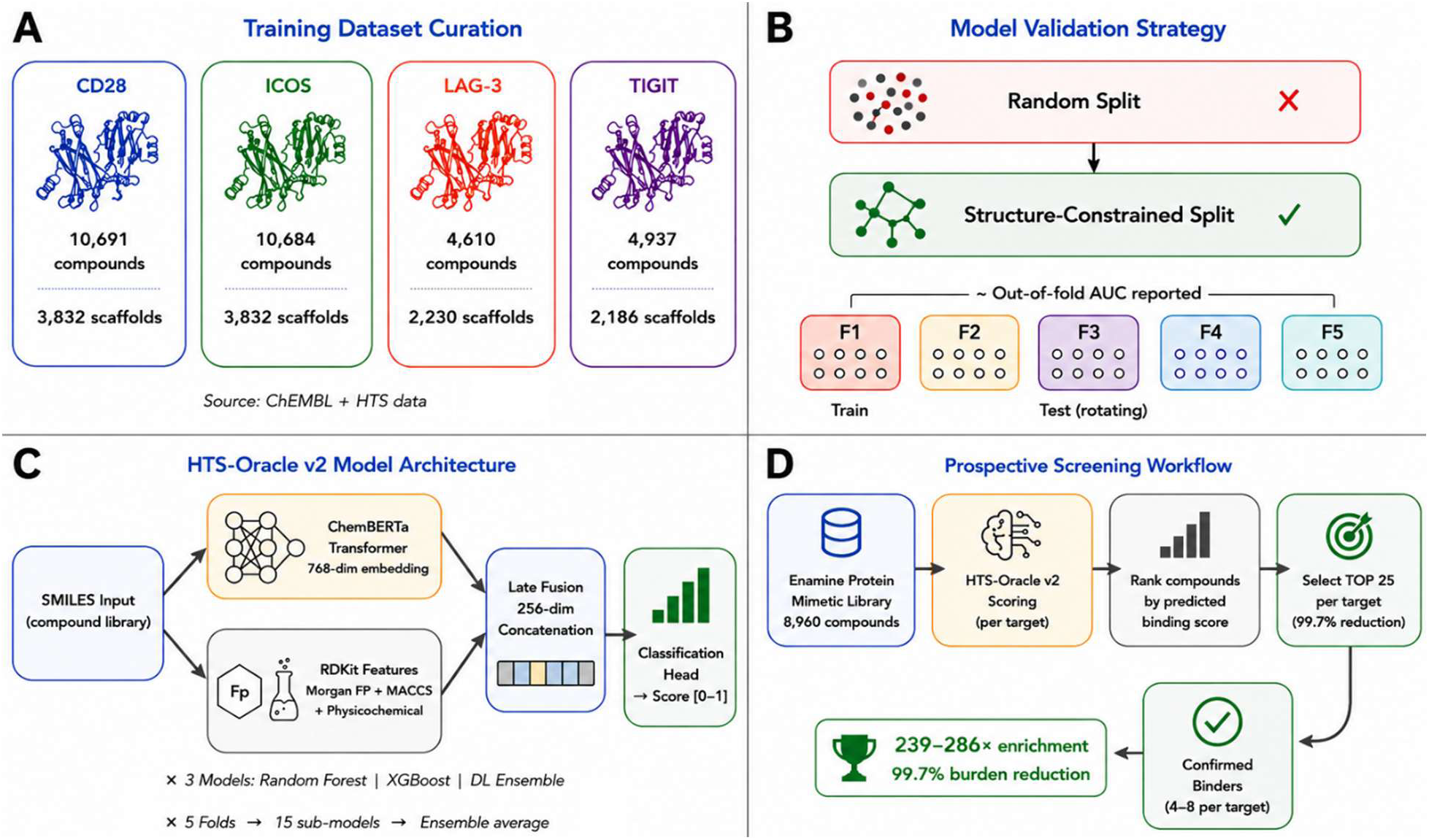
Overview of HTS-Oracle v2 and the prospective discovery workflow for functional small molecule modulators. **(A)** Training dataset composition for CD28, ICOS, LAG-3, and TIGIT, including the number of compounds and unique scaffolds curated from ChEMBL and HTS datasets. **(B)** Model validation strategy comparing random split and structure-constrained split approaches used during model development prior to prospective deployment. **(C)** HTS-Oracle v2 architecture integrating ChemBERTa embeddings and molecular descriptors through ensemble machine learning models for compound prioritization. **(D)** Prospective screening workflow applied to the 8,960-compound Enamine Protein Mimetic Library, including model-based ranking, experimental validation, and identification of functional small molecule modulators.

HTS-Oracle v2 features the implementation of rigorous cross-validation throughout all model development (Figure 1B). Compounds sharing the same core scaffold were assigned exclusively to the same fold, ensuring zero scaffold leakage between training and test sets across all five iterations. This directly addresses a well-documented limitation of random splits in virtual screening benchmarking, where structural similarity between training and test compounds artificially inflates performance estimates and misrepresents prospective generalizability. The cross-validation strategy produced realistic fold imbalance in the distribution of known actives across individual test folds, ranging from 4.8% to 28.7% depending on the target (Figure S1). These percentages reflect the concentration of curated known binders within each scaffold-partitioned fold, not experimental HTS hit rates, which were substantially lower (0.056–0.089% as established by full-library random screening), and confirm that the splits are genuinely scaffold-diverse rather than artificially balanced. This deliberate structural diversity between folds provides performance estimates that honestly reflect what a model would encounter when applied to a new, structurally distinct compound library. HTS-Oracle v2 employs a multimodal ensemble architecture combining ChemBERTa transformer embeddings with RDKit-derived cheminformatics descriptors across three parallel model classes, trained under this rigorous framework (Figure 1C). The trained models were subsequently deployed prospectively against the 8,960-compound Enamine Protein Mimetic Library, with compound selection, experimental screening, and hit validation conducted as described in the workflow overview (Figure 1D).

### Computational Model Performance Across Four Immune Checkpoint Targets

HTS-Oracle v2 was trained using three complementary model architectures, Random Forest (RF), XGBoost (XGB), and a deep learning (DL) ensemble combining ChemBERTa transformer embeddings with RDKit-derived cheminformatics descriptors, across all four targets under rigorous cross-validation. Full performance metrics are summarized in Table 1 and Figure S2.

**Table 1.**
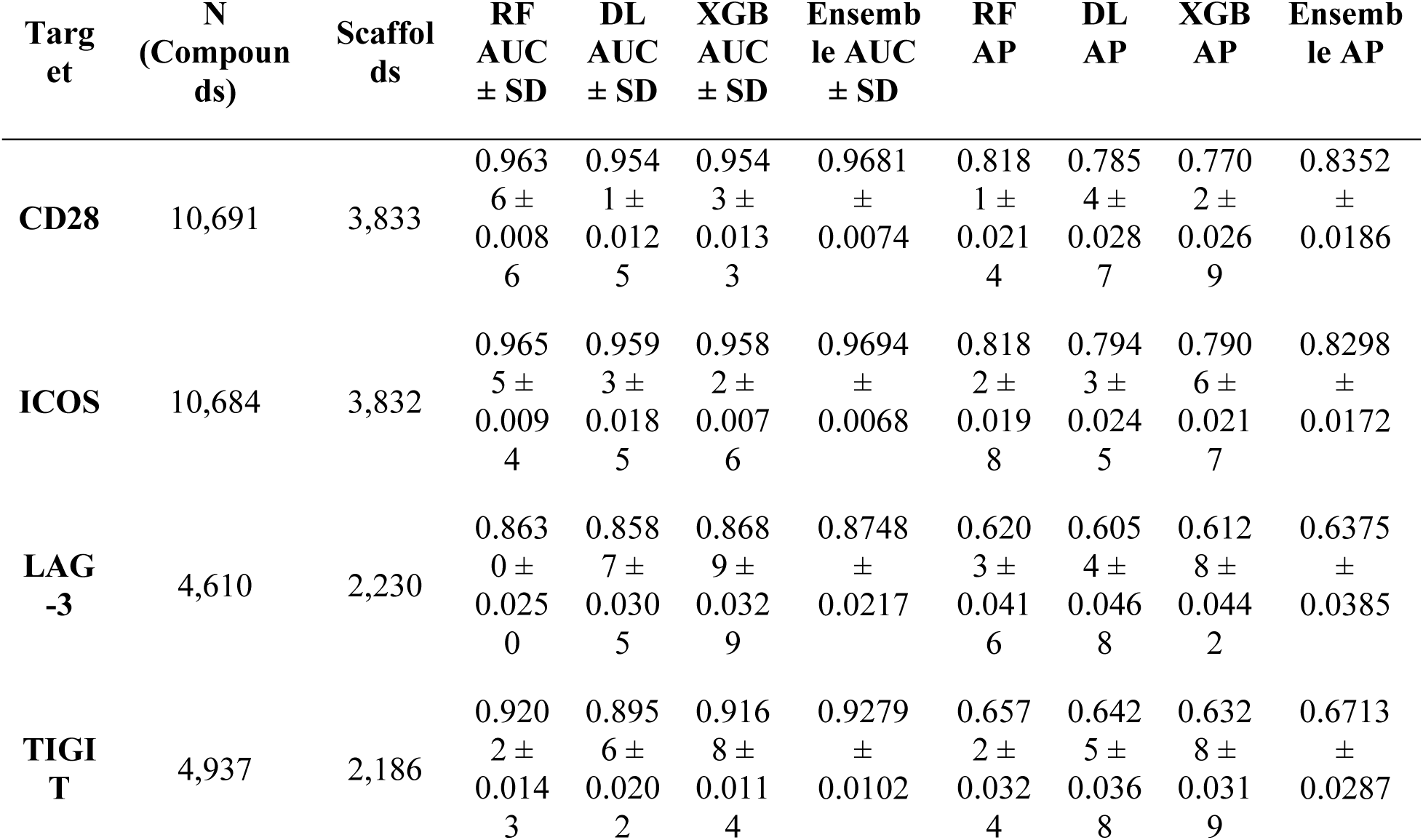
Model performance metrics for HTS-Oracle v2 across four immune checkpoint targets under rigorous cross-validation. Reported metrics include ROC-AUC ± SD and average precision (AP) ± SD for the individual Random Forest (RF), XGBoost (XGB), and Deep Learning (DL) models, as well as the final ensemble model used for prospective compound prioritization and experimental validation.

For CD28 and ICOS, all three model architectures achieved strong discriminative performance, with RF ROC-AUC values of 0.9636 ± 0.0086 and 0.9655 ± 0.0094 respectively (Table 1), supported by average precision (AP) values of 0.818 for both targets. The consistency of performance across all five scaffold-aware folds for these targets, CD28: 0.953–0.978; ICOS: 0.953–0.982 (Figure S2), demonstrates robust learning of structure-activity relationships despite rigorous structural separation between training and test sets. For TIGIT, RF achieved an AUC of 0.9202 ± 0.0143, representing strong performance for a target with approximately half the training data of CD28 and ICOS. LAG-3 showed the most modest performance with RF AUC of 0.8630 ± 0.0250 (Table 1), consistent with its smallest dataset size and highest scaffold diversity ratio, indicative of a chemically diverse active set that is inherently more challenging to generalize across scaffold boundaries. Critically, all four models remain well above the AUC threshold considered useful for virtual screening applications (AUC > 0.7), confirming that HTS-Oracle v2 provides meaningful compound prioritization across the full target panel under rigorous validation conditions (Figure 2).

**Figure 2.**
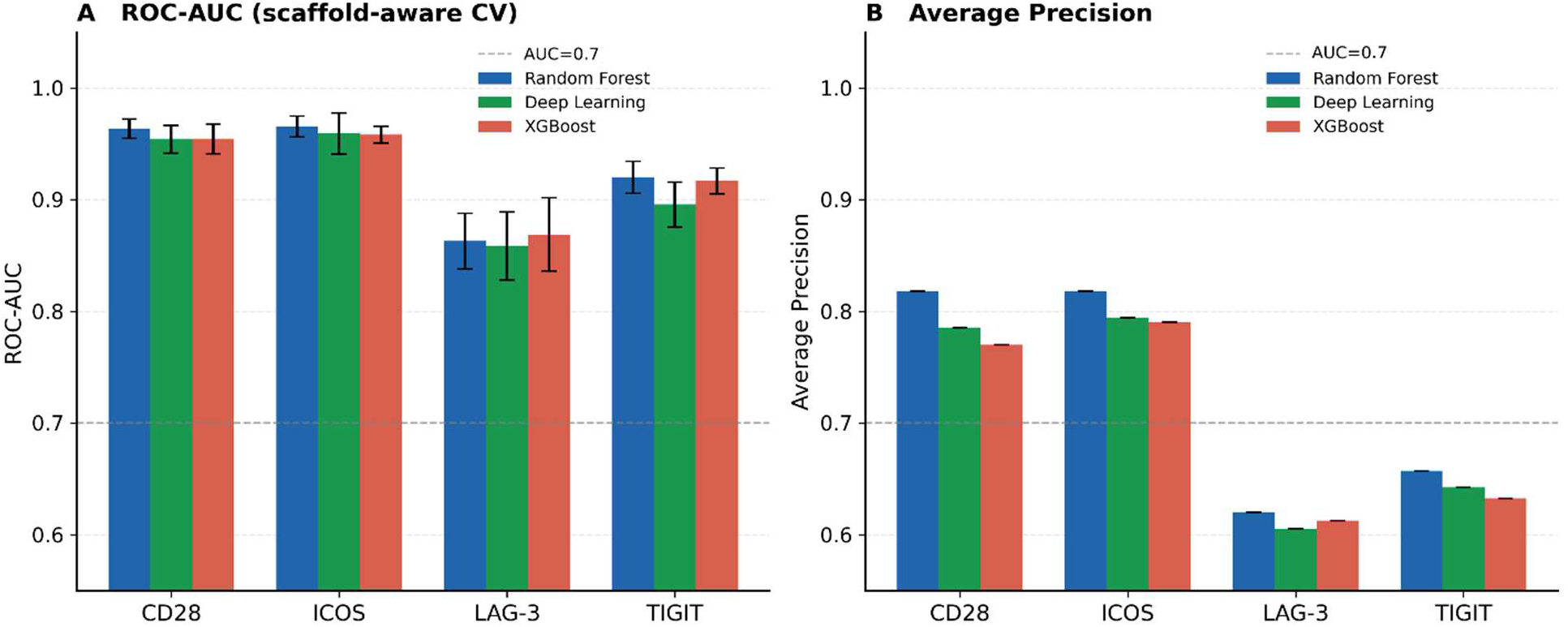
HTS-Oracle v2 model performance across four immune checkpoint targets under rigorous cross-validation. **(A)** ROC-AUC values for Random Forest (RF), Deep Learning (DL) ensemble, and XGBoost (XGB), reported as mean ± SD across five scaffold-aware folds. The dashed line indicates the AUC = 0.7 threshold considered useful for virtual screening. **(B)** Average precision (AP) for all three model architectures, reflecting early enrichment performance under class imbalance conditions. All models were evaluated using out-of-fold predictions aggregated across all five scaffold-aware folds.

Across all four targets, RF consistently matched or outperformed XGBoost and the DL ensemble under scaffold-aware CV (Table 1, Figure 2), with the most pronounced gap observed for TIGIT (RF: 0.920 vs DL: 0.896). This pattern is consistent with the known behavior of tree ensemble methods under data-scarce, high-diversity conditions, where the inductive bias of random forests toward individual feature importance confers more robust generalization than the more flexible but data-hungry deep learning architectures. The DL ensemble remained competitive within approximately 0.01 AUC of RF across all targets (Figure S2), demonstrating that the multimodal architecture provides consistent value even if it does not uniformly surpass simpler baselines under scaffold-aware validation.

Average precision values, which are more informative than AUC under conditions of class imbalance, showed a consistent pattern mirroring the AUC results (Table 1): CD28 and ICOS achieved RF AP of 0.818, substantially higher than LAG-3 (AP = 0.620) and TIGIT (AP = 0.657) (Figure 2B). This gradient directly reflects the dependence of model quality on training data volume, a finding with important practical implications for target selection in AI-guided drug discovery programs. Specifically, targets with larger, more structurally diverse training datasets consistently yield models with superior early enrichment properties, as captured by average precision, beyond what AUC alone would suggest. The ensemble framework (Table 1) consistently matched or outperformed the individual RF, XGB, and DL models across all four targets, indicating that integrating complementary model predictions provided a robust strategy for prospective hit prioritization.

### Full Library Random Screening Establishes Rigorous Experimental Baselines

To establish a rigorous experimental baseline for enrichment factor calculation, all 8,960 compounds of the Enamine Protein Mimetic Library were subjected to full-library single-dose screening against CD28, ICOS, LAG-3, and TIGIT using the Dianthus NT.23 Pico instrument with TRIC technology (Figure 3). Compounds exhibiting ΔF_norm_ values greater than three standard deviations from the negative control were designated as primary hits. Compounds exhibiting autofluorescence, quenching, or aggregation artifacts were identified through orthogonal control experiments and excluded prior to confirmation.

**Figure 3.**
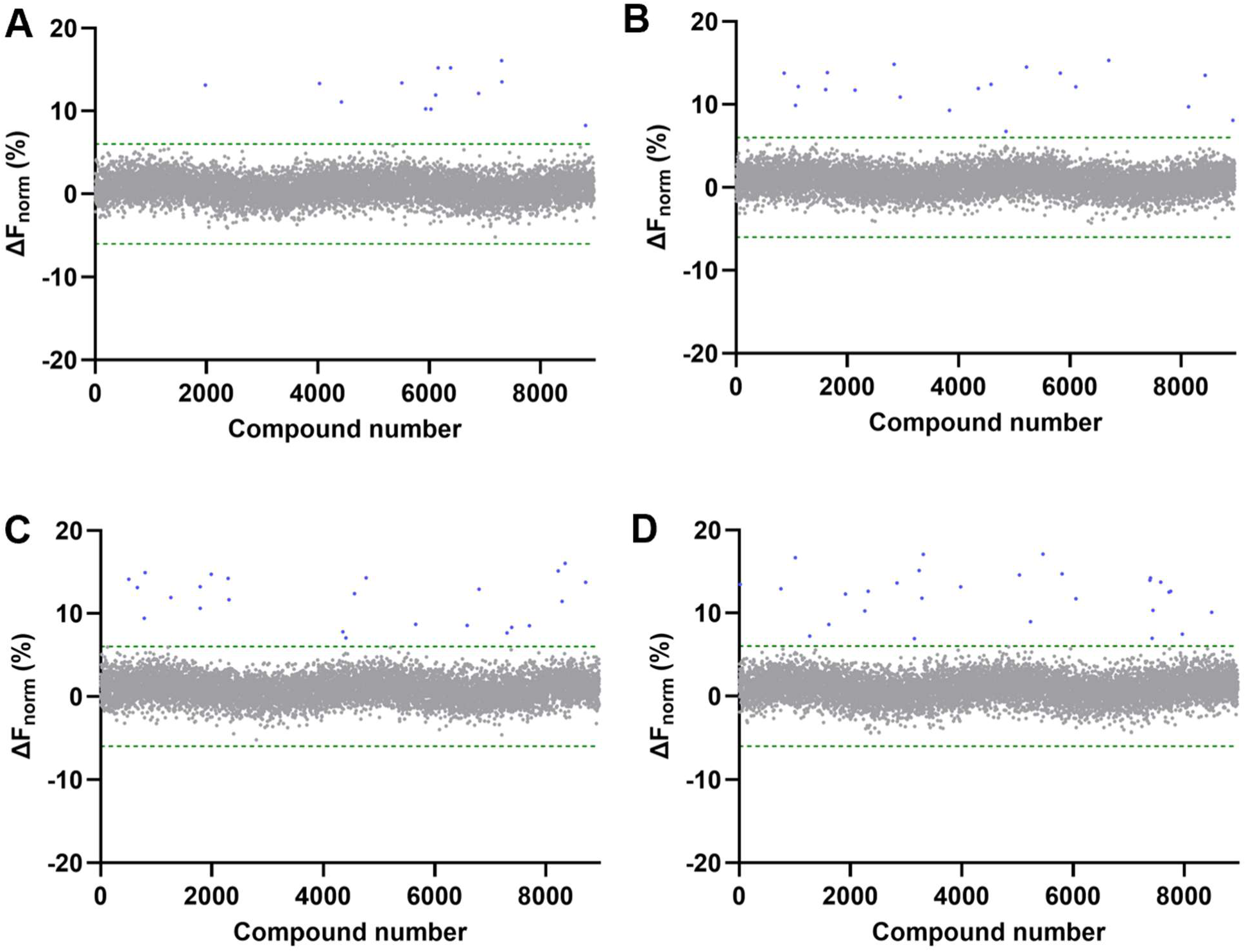
Full-library single-dose TRIC screening of all 8,960 Enamine Protein Mimetic Library compounds against **(A)** CD28, **(B)** ICOS, **(C)** LAG-3, and **(D)** TIGIT. ΔF_norm_ values are plotted for each compound. Gray dots represent non-binders; blue dots represent primary hits exceeding the hit threshold; green dashed lines indicate the mean ± 3 SD of negative controls. Primary hit rates were 0.145%, 0.212%, 0.268%, and 0.312% for CD28, ICOS, LAG-3, and TIGIT respectively.

Full-library random screening identified primary hit rates of 0.145%, 0.212%, 0.268%, and 0.312% for CD28, ICOS, LAG-3, and TIGIT respectively across the 8,960-compound library (Figure 3, Table S1). All primary hits were subsequently subjected to dose-response binding affinity measurements using Monolith X spectral shift detection to determine dissociation constants and exclude false positives arising from nonspecific or artifactual binding (Table S1). Following dose-response validation, confirmed validated hit rates were 0.067%, 0.078%, 0.056%, and 0.089% for CD28, ICOS, LAG-3, and TIGIT respectively (Table S1), reflecting the inherent difficulty of identifying small molecule binders for these PPI-driven proteins. The complete list of all validated hits across all four targets, including compound structures, SMILES, and binding affinities, is provided in Table S2. The attrition rate from primary to validated hits ranged from 46% for CD28 to 79% for LAG-3. These rigorously established baseline values were used as denominators for all subsequent enrichment factor calculations, ensuring that enrichment reflects performance against a stringent experimental ground truth.

### Prospective Validation of HTS-Oracle v2: Improved Enrichment Relative to Random Selection Across Four Immune Checkpoint Targets

HTS-Oracle v2 was applied prospectively to the full 8,960-compound Enamine Protein Mimetic Library, generating per-target ensemble binding probability scores derived from the RF, XGB, and DL models. Compounds were ranked by their target-specific model score, and the top 25 compounds per target were selected for experimental validation, representing a 99.7% reduction in screening burden relative to full-library testing (Tables S3-S6). Experimental validation followed identical protocols to the random screening baseline, including single-dose TRIC screening, orthogonal artifact exclusion, and dose-response confirmation for all primary hits (Figure 4). To provide additional structural context, the maximum Tanimoto similarity between each experimentally validated hit and compounds present in the corresponding target-specific training dataset was calculated using Morgan fingerprints. Similarity values ranged from 0.48 to 0.72 across validated hits (Table S7), indicating that the identified compounds were related to, but not identical to, previously observed chemotypes.

**Figure 4.**
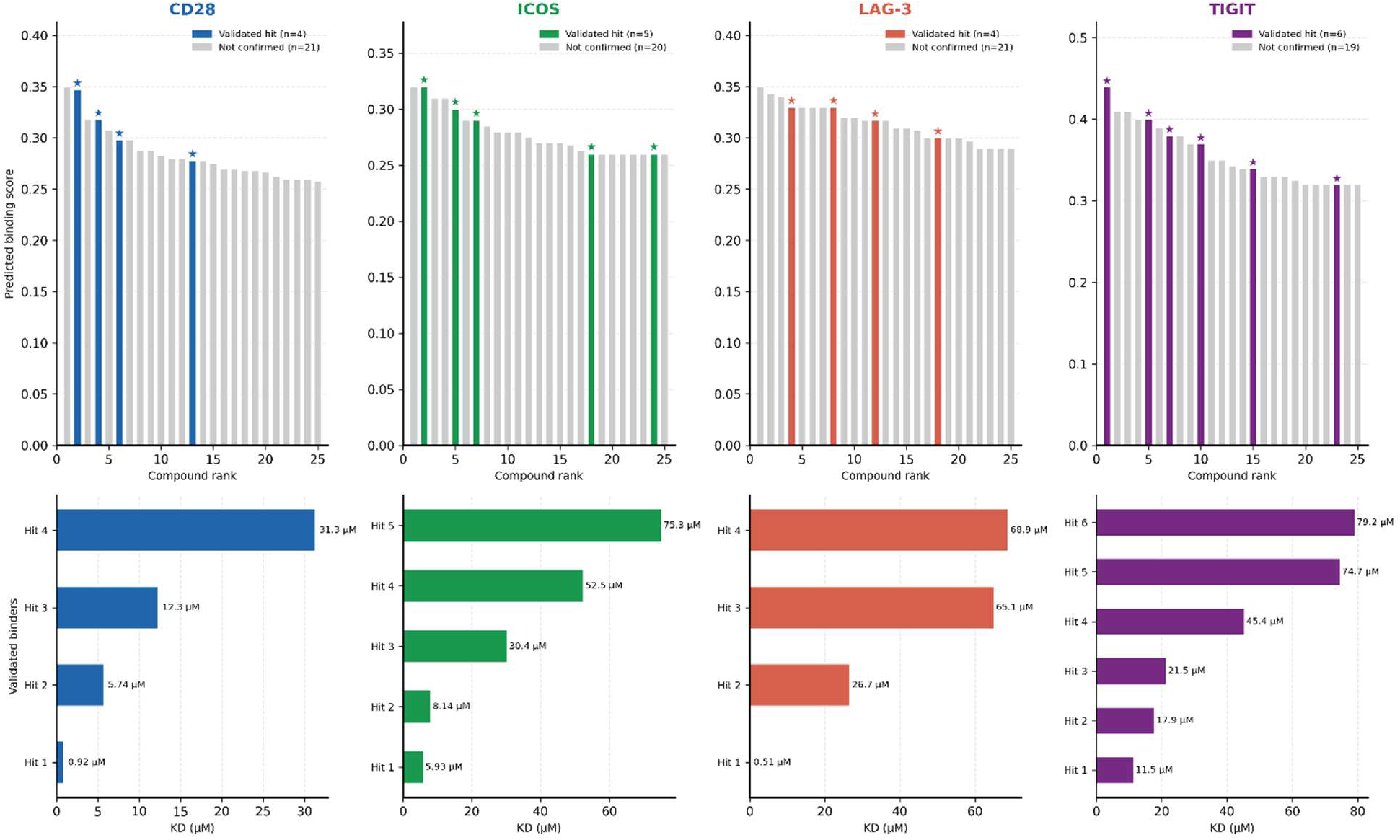
HTS-Oracle v2 prospective screening results for the top 25 model-selected compounds per target. Upper panels show predicted binding scores for all 25 ranked compounds, with validated hits highlighted in color (★) and non-confirmed compounds shown in gray. Lower panels show KD values for all confirmed binders per target, determined by Monolith X spectral shift detection.

HTS-Oracle v2 yielded substantial prospective enrichment across all four targets relative to random selection (Table 2, Figure 5). For CD28, 4 of 25 model-selected compounds were confirmed as validated binders, corresponding to a validated hit rate of 16.0% and an enrichment factor of 239× over the full-library baseline validated hit rate of 0.067% (Table 2). For ICOS, 5 of 25 compounds were confirmed, yielding a 20.0% hit rate and 256× enrichment over a 0.078% baseline. For LAG-3, 4 validated hits were identified among 25 model-selected compounds, giving a 16.0% hit rate and the highest single-target enrichment factor of 286× over a 0.056% baseline. For TIGIT, 6 of 25 compounds were confirmed, corresponding to a 24.0% hit rate and 270× enrichment over a 0.089% baseline. These results are summarized in Table 2. Binding affinities of all confirmed HTS-Oracle v2 hits are reported in Table S2.

**Figure 5.**
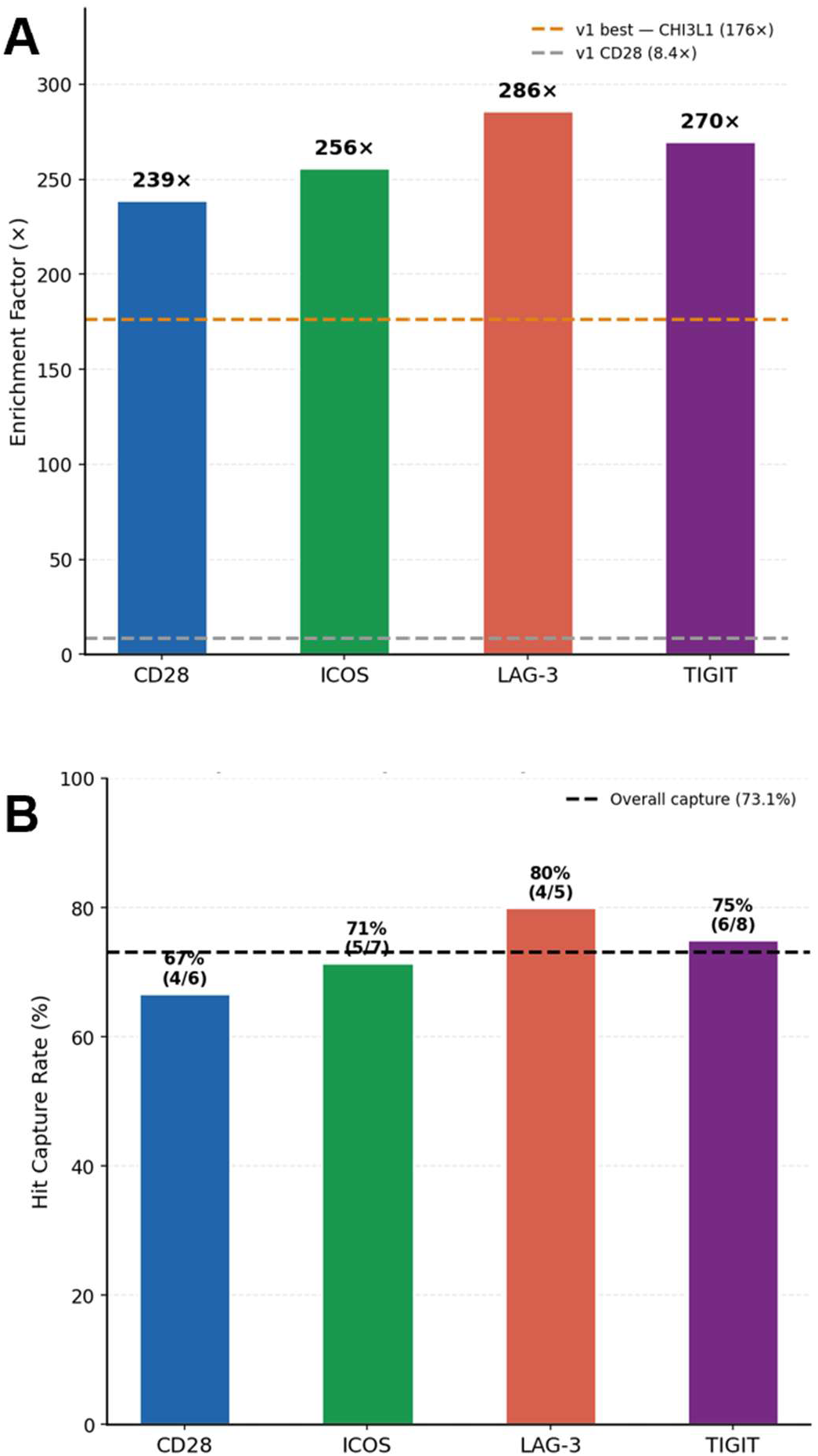
HTS-Oracle v2 prospective validation performance across four immune checkpoint targets. **(A)** Enrichment factors relative to full-library random screening validated hit rates. Dashed lines indicate HTS-Oracle v1 performance benchmarks for CHI3L1 (176×, best result across all v1 targets) and CD28 (8.4×, the only target shared between v1 and v2). All four HTS-Oracle v2 targets exceed both v1 benchmarks. **(B)** Hit capture rates showing the percentage of all experimentally validated library hits recovered within the top 25 model-selected compounds per target. The dashed line indicates the overall hit capture rate of 73.1% across all four targets combined.

**Table 2.** HTS-Oracle v2 prospective validation results across four immune checkpoint targets. Compounds tested refers to the top 25 model-selected compounds per target from the 8,960-compound Enamine Protein Mimetic Library. Validated hit rate was determined after dose-response confirmation by Monolith X spectral shift detection. Enrichment factor was calculated relative to the full-library random screening validated hit rate. Hit capture rate represents the percentage of all validated hits identified in full-library random screening that were recovered within the top 25 model-selected compounds. The overall row summarizes performance across all four targets combined.

| Target | Compounds tested | Validated hits | Hit rate % | Baseline rate % | Enrichment factor (x) | Library hits captured | Capture rate (%) |
| --- | --- | --- | --- | --- | --- | --- | --- |
| **CD28** | 25 | 4 | 16.0% | 0.067% | 239x | 4/6 | 66.7% |
| **ICOS** | 25 | 5 | 20.0% | 0.078% | 256x | 5/7 | 71.4% |
| **LAG-3** | 25 | 4 | 16.0% | 0.056% | 286x | 4/5 | 80.0% |
| **TIGIT** | 25 | 6 | 24.0% | 0.089% | 270x | 6/8 | 75.0% |

A particularly striking finding is the hit capture rate achieved by HTS-Oracle v2 (Figure 5B). Of the 26 total validated hits identified across the entire 8,960-compound library by random screening, 19 were captured within the 100 model-selected compounds tested across all four targets, an overall hit capture rate of 73.1%. On a per-target basis, hit capture rates were 66.7% for CD28 (4 of 6 library hits), 71.4% for ICOS (5 of 7), 80.0% for LAG-3 (4 of 5), and 75.0% for TIGIT (6 of 8) (Figure 5B). This means HTS-Oracle v2 recovered approximately three-quarters of the library’s true experimentally confirmed binders by testing just 0.3% of the compound collection per target, a result that directly demonstrates the model’s capacity to concentrate genuine binding activity at the top of its ranked list (Figure S4).

The consistency of enrichment across all four targets, spanning both costimulatory (CD28, ICOS) and co-inhibitory (LAG-3, TIGIT) immune checkpoint axes, and across both large and small training datasets, demonstrates the generalizability of the training approach across diverse immune checkpoint biology (Figure 5A). Furthermore, the strong performance observed for LAG-3 (286× enrichment relative to random selection) despite its lower retrospective AUC (0.863) relative to CD28 and ICOS highlights an important and underappreciated principle: prospective enrichment in a real screening campaign is determined not only by model AUC but also by the background hit rate of the target library. Against the extremely low baseline hit rate of 0.056% for LAG-3, even a model with modest discriminative power can achieve practical enrichment if it correctly ranks even a small number of genuine binders near the top of the list.

The most potent binder identified per target, Z332849278 for CD28 (KD = 0.92 ± 0.08 µM), Z338150202 for ICOS (KD = 5.93 ± 0.62 µM), Z1282765290 for LAG-3 (KD = 0.51 ± 0.02 µM), and Z108564456 for TIGIT (KD = 11.5 ± 1.14 µM), demonstrated dose-dependent, saturable binding isotherms with well-defined dissociation constants, confirming specific and high-quality target engagement (Figure 6A-H). Notably, the top LAG-3 and CD28 binders achieved sub-micromolar and low single-digit micromolar affinities respectively, representing among the most potent small molecule binders reported for these immune checkpoint targets to date. To confirm that the identified binders functionally disrupt their respective immune checkpoint protein–protein interactions, the most potent hit per target was evaluated in orthogonal functional assays. CD28 binding was assessed using a competitive CD28/CD80 ELISA, ICOS using an ICOS/ICOSL ELISA, LAG-3 using a LAG-3/MHC-II FRET assay, and TIGIT using a TIGIT/CD155 AlphaLISA. All four top binders demonstrated dose-dependent inhibition of their respective protein-protein interactions, with IC_50_ values of 1.41 ± 0.07 µM for CD28 (Figure 6I), 4.23 ± 0.51 µM for ICOS (Figure 6J), 0.36 ± 0.07 µM for LAG-3 (Figure 6K), and 8.67 ± 0.92 µM for TIGIT (Figure 6L), confirming on-target functional activity and validating HTS-Oracle v2’s capacity to identify not only binding compounds but functionally relevant inhibitors of immune checkpoint signaling.

**Figure 6.**
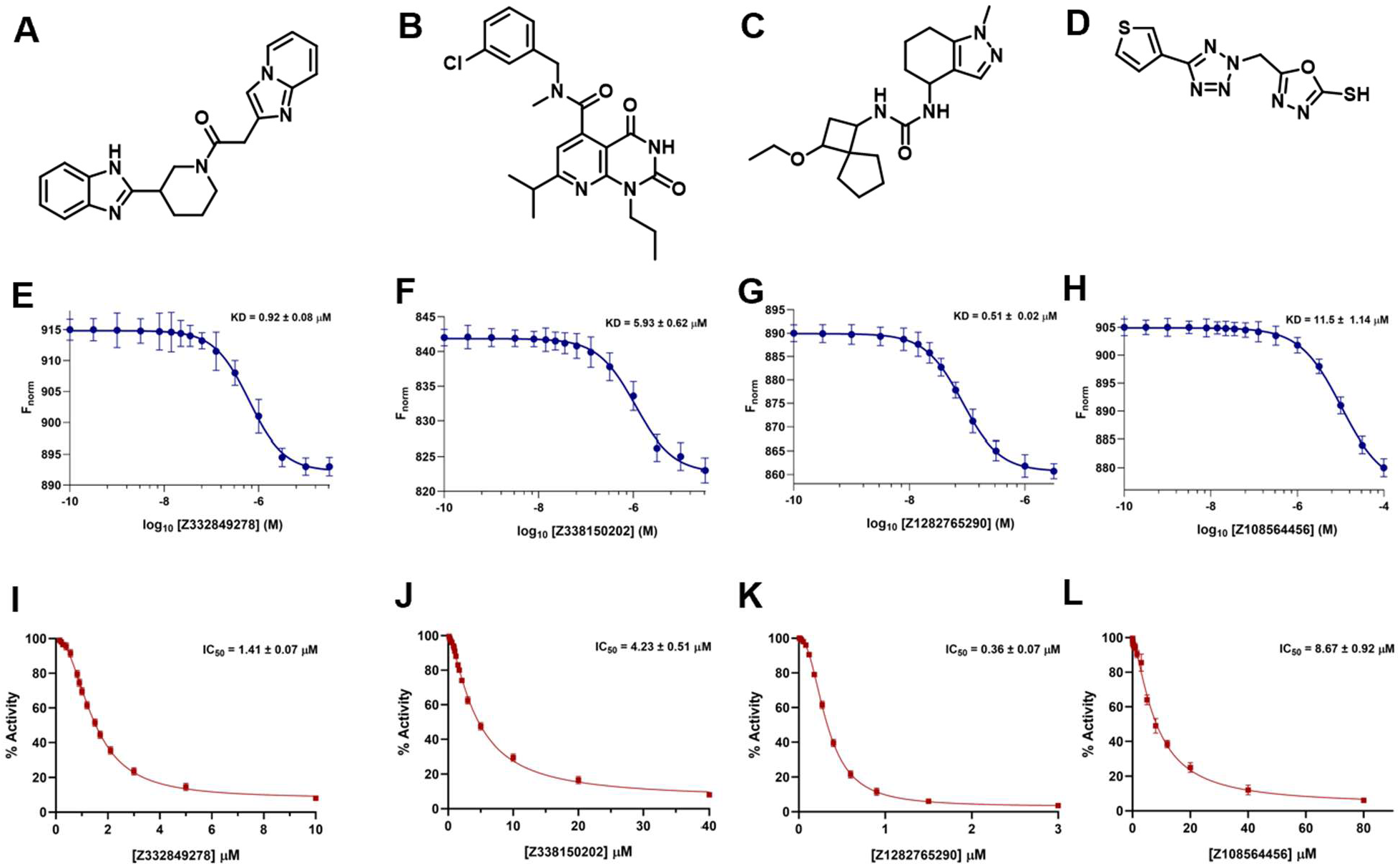
Binding affinity and functional blockade of the top HTS-Oracle v2-predicted hit per immune checkpoint target. **A-D** panels display the top hit compound per CD28, ICOS, LAG-3, and TIGIT, respectively. **E-H** panels show dose-response binding isotherms determined by Monolith X spectral shift detection. **I-L** panels show dose-dependent inhibition of protein-protein interactions determined by CD28/CD80 competitive ELISA, ICOS/ICOSL ELISA, LAG-3/MHC-II FRET assay, and TIGIT/CD155 AlphaLISA for CD28, ICOS, LAG-3, and TIGIT respectively. IC_50_ values were determined by four-parameter nonlinear regression using GraphPad Prism 10. Data represent mean ± SD from three independent experiments.

From a medicinal chemistry perspective, the top validated compounds identified across the four targets represent structurally diverse chemotypes (Figure 6). The lead CD28 modulator Z332849278 is built around a fused bicyclic heteroaromatic core, whereas the top ICOS hit Z338150202 contains a distinct substituted aromatic heterocycle scaffold. The most potent LAG-3 modulator, Z1282765290, incorporates a chemically differentiated nitrogen-rich heterocyclic framework, while the leading TIGIT compound Z108564456 features a separate aromatic heterocycle architecture. Collectively, these compounds span multiple scaffold classes and demonstrate that HTS-Oracle v2 can identify structurally diverse, experimentally validated modulators rather than repeatedly prioritizing a single chemotype. These findings provide multiple chemically distinct starting points for future medicinal chemistry optimization.

### HTS-Oracle v2 Outperforms v1 for CD28. A Direct Head-to-Head Comparison

CD28 represents the critical bridge between HTS-Oracle v1 and v2, as the only target shared between both generations of the platform and historically the most challenging case for the first-generation model. In HTS-Oracle v1, CD28 model training used a severely imbalanced dataset with a ∼2% active rate under random cross-validation splits, and prospective screening of 345 model-selected compounds from a 1,152-compound in-house library yielded an 8.4-fold enrichment over the ∼1% random HTS baseline, the lowest enrichment achieved across any target in v1, alongside a 70% reduction in screening burden.^27^

Training on a substantially expanded CD28 dataset in HTS-Oracle v2 under rigorous CV produced an AUC of 0.9681 ± 0.0074, a more rigorous and honest performance estimate than v1’s random-split metrics. Prospectively, HTS-Oracle v2 identified 4 validated CD28 binders among just 25 model-selected compounds from the 8,960-compound Enamine library, yielding 239× enrichment (relative to random selection) compared to 8.4× for v1 (Figure 7A) and a 16.0% validated hit rate compared to 8.41% for v1 (Figure 7B), despite v1 testing 345 compounds versus only 25 for v2. This represents a 28-fold improvement in enrichment factor alongside a step change in screening burden reduction from 70% to 99.7% (Figure 7C). The improvement reflects two compounding factors: first, the larger and more balanced training dataset provides richer structural coverage of the CD28 binding pharmacophore; second, rigorous CV during training enforces generalization to novel scaffolds, directly improving the model’s ability to correctly rank previously unseen compounds. The improved performance gain on CD28, the target where v1 struggled most, provides the strongest evidence that the advances in HTS-Oracle v2 translate directly into improved prospective performance.

**Figure 7.**
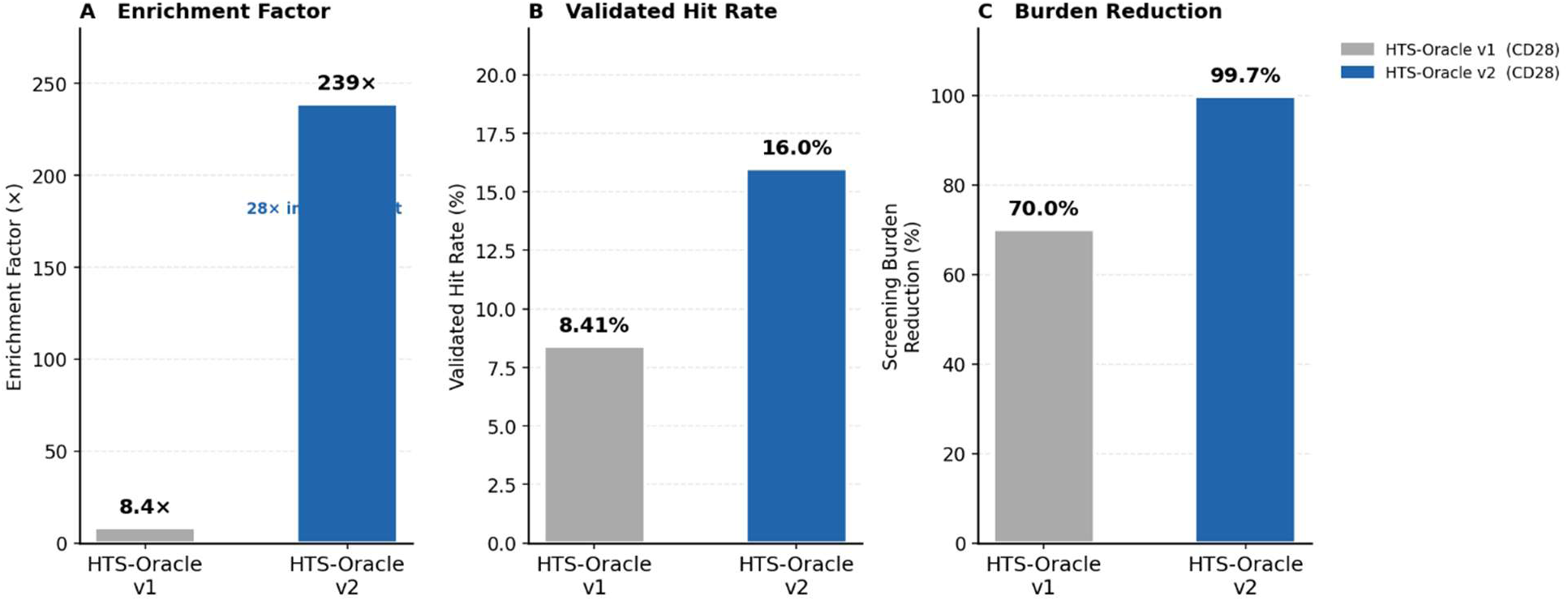
Direct head-to-head comparison of HTS-Oracle v1 and HTS-Oracle v2 for CD28. **(A)** Enrichment factor achieved by each platform relative to experimental screening baselines. **(B)** Validated hit rate obtained during prospective screening. **(C)** Screening burden reduction achieved by each platform. **(D)** Schematic comparison of training dataset composition and methodological differences between HTS-Oracle v1 and HTS-Oracle v2.

### Relationship Between Training Data Volume and Model Performance

A consistent and interpretable pattern emerges across the four-target panel: model performance under scaffold-aware CV is strongly and positively correlated with training dataset size (Figure 8). CD28 and ICOS, each with approximately 10,700 compounds, achieve RF AUC values of 0.964 and 0.966 respectively. TIGIT, with approximately 4,900 compounds, achieves an intermediate AUC of 0.920. LAG-3, the smallest dataset at approximately 4,600 compounds, shows the most modest AUC of 0.863 with the highest per-fold variance, reflecting greater sensitivity to scaffold-specific fold composition under data-scarce conditions.

**Figure 8.**
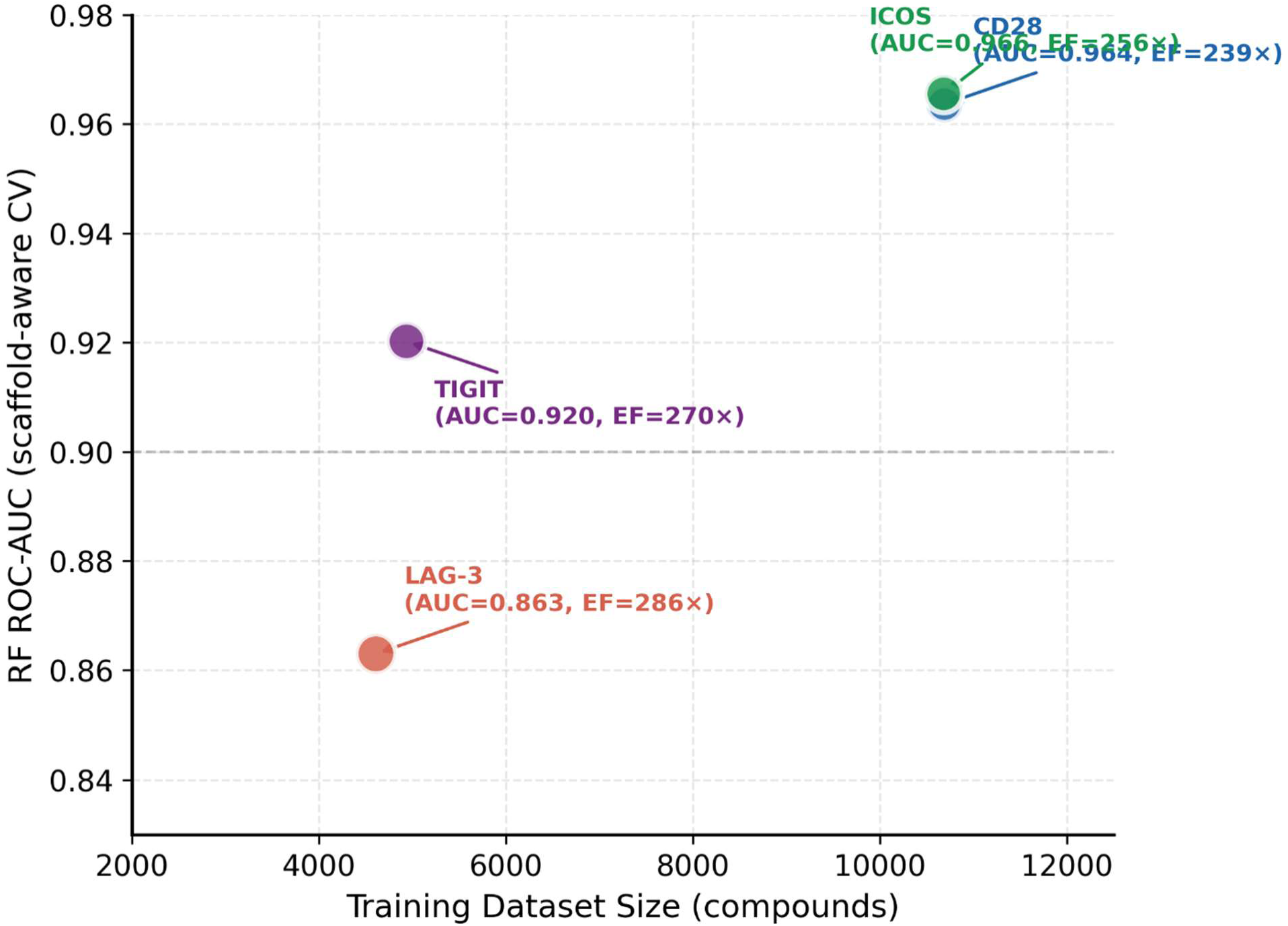
Relationship between training dataset size, scaffold-aware cross-validation performance, and prospective enrichment factor across four immune checkpoint targets. Each data point represents one target; bubble size is proportional to the prospective enrichment factor achieved in experimental validation. The positive correlation between dataset size and ROC-AUC confirms the data-volume dependence of model quality under scaffold-aware validation. Notably, LAG-3 achieves the highest enrichment factor (286×) despite its lowest AUC, reflecting the dominant contribution of the extremely low background hit rate (0.056%) to prospective enrichment performance.

Importantly, this data-size dependence does not translate linearly into prospective performance (Figure 8). LAG-3 achieved the highest single-target enrichment factor (286×) relative to random selection despite having the lowest AUC, because its extremely low baseline hit rate (0.056%) means that even modest discriminative power at the top of the ranked list translates into exceptional enrichment. This decoupling of retrospective AUC from prospective enrichment underscores a critical practical principle: AUC alone is insufficient to predict the real-world utility of a virtual screening model, and the background hit rate of the target library is an equally important determinant of prospective value. These observations suggest that for future AI-guided screening campaigns against immune checkpoint targets, dataset expansion through active learning, transfer learning, or incorporation of negative screening data should be prioritized for data-scarce targets such as LAG-3 and TIGIT to further improve both retrospective and prospective performance.

## Conclusions

In this study, we demonstrate the prospective application of HTS-Oracle v2 for the discovery and experimental validation of functional small molecule immune checkpoint modulators across four therapeutically relevant targets. Trained and validated simultaneously across four clinically significant immune checkpoint targets, CD28, ICOS, LAG-3, and TIGIT, HTS-Oracle v2 achieves ROC-AUC values of 0.875-0.969 under rigorous cross-validation, demonstrating robust and consistent discriminative performance across diverse chemical space and varying training dataset sizes.

Prospective experimental validation against an 8,960-compound Enamine Protein Mimetic Library confirmed the platform’s real-world utility in unambiguous terms. Selecting just 25 compounds per target, a 99.7% reduction in screening burden, HTS-Oracle v2 recovered 67–80% of all experimentally validated library hits with enrichment factors of 239–286× over rigorously established full-library random screening baselines. For CD28, the target shared with HTS-Oracle v1, v2 delivers a 28-fold improvement in enrichment factor and a step change in screening burden reduction from 70% to 99.7%, providing direct experimental evidence that rigorous validation translates into superior prospective performance. Furthermore, the top binders identified per target demonstrated potent, dose-dependent binding affinities and functionally disrupted their respective immune checkpoint protein–protein interactions, confirming that HTS-Oracle v2 enriches not only for binding compounds but for functionally relevant inhibitors of immune checkpoint signaling.

We anticipate that its application to additional immune checkpoint targets, combined with continued dataset expansion and integration of functional screening data, will further extend its utility across the broader landscape of PPI-driven immune modulators, targets that remain among the most clinically important yet pharmacologically underexplored in modern drug discovery.

## Material and Methods

### Dataset Curation and Preparation

Training compounds were curated from ChEMBL and high-throughput screening datasets containing target-specific activity annotations for CD28, ICOS, LAG-3, and TIGIT. The resulting datasets exhibited active-to-inactive ratios of approximately 1:20, 1:18, 1:22, and 1:17 for CD28, ICOS, LAG-3, and TIGIT, respectively. Because the datasets were assembled from multiple assay formats and literature sources, activity labels were assigned according to the classifications reported in the original datasets rather than by applying a single uniform activity threshold across all compounds. Each compound was assigned a binary activity label, where y = 1 if its identifier appeared in the confirmed actives set and y = 0 otherwise. SMILES strings were validated using RDKit parsing prior to featurization, with error-handling fallbacks applied to ensure pipeline integrity. Invalid SMILES strings were excluded from analysis. All datasets were curated to remove duplicate entries based on compound identifier, and canonical SMILES were generated for all retained compounds using RDKit standardization procedures.

### Molecular Featurization

Each compound was represented using three complementary molecular feature sets, following the approach established in HTS-Oracle v1.^27^ Morgan circular fingerprints were computed with radius 2 and 2,048 bits, encoding circular neighborhood substructures around each atom. In parallel, 167-bit MACCS structural keys were generated to capture common pharmacophoric substructural motifs. Additionally, 15 physicochemical descriptors were computed using RDKit, including molecular weight, LogP, topological polar surface area (TPSA), hydrogen bond donors, hydrogen bond acceptors, rotatable bonds, ring count, aromatic ring count, fraction of sp3 carbons, number of heteroatoms, and quantitative estimate of drug-likeness (QED). These three feature vectors were concatenated to form a 2,230-dimensional feature vector per compound. All descriptor values were sanitized by replacing NaN and infinite values with zeros prior to model training, and features were standardized using StandardScaler to normalize scales across descriptors.

### Cross-Validation

Murcko scaffolds were extracted for all compounds in each dataset using RDKit’s MurckoDecompose function. Compounds sharing identical Murcko scaffolds were assigned exclusively to the same fold, ensuring complete structural separation between training and test sets across all five cross-validation iterations. This scaffold-based partitioning prevents molecular leakage, the phenomenon whereby structurally similar molecules appear in both training and test sets under random splitting, artificially inflating performance estimates and misrepresenting prospective generalizability. Fold balance in terms of active compound percentage was not artificially enforced, allowing the natural distribution of actives across scaffold clusters to determine fold composition. This produced realistic variation in active rates across folds, ranging from 4.8% to 28.7% depending on the target and fold, reflecting the genuine structural heterogeneity of the training datasets.

### Model Development

Three model architectures were trained and evaluated under scaffold-aware cross-validation for each target.

#### Random Forest

Random Forest classifiers were trained using scikit-learn with 200 estimators, the Gini impurity criterion, and default hyperparameters otherwise. Class imbalance was addressed through balanced class weights computed from the training fold class distribution at each cross-validation iteration. Out-of-fold predictions were aggregated across all five folds to generate global ROC-AUC and average precision metrics.

#### XGBoost

XGBoost classifiers were trained with 200 estimators and a scale_pos_weight parameter set to the ratio of negative to positive examples in each training fold, providing explicit correction for class imbalance. All other hyperparameters were set to XGBoost defaults.

#### Deep Learning Ensemble

The deep learning component follows the multimodal ensemble architecture described in HTS-Oracle v1,^27^ comprising two parallel processing branches that merge via late fusion. The first branch employs ChemBERTa (seyonec/ChemBERTa-zinc-base-v1),[14] a RoBERTa-based transformer pretrained on SMILES strings, to generate 768-dimensional contextual molecular embeddings from the classification [CLS] token. These embeddings are regularized through dropout and processed through a multilayer perceptron (MLP) with architecture 768→256→128. The second branch processes the 2,230-dimensional cheminformatics feature vector through three parallel feature selection strategies, LASSO regression, principal component analysis (PCA), and mutual information filtering, each reducing the feature space to 200 components, followed by a parallel MLP of identical architecture. The 128-dimensional outputs from both branches are concatenated into a 256-dimensional representation and passed through a final classification head (256→128→64→1) with batch normalization, ReLU activations, and graduated dropout. The ensemble trains 15 sub-models, one per feature selection method per cross-validation fold, and averages predictions across all sub-models to produce the final ensemble score. Training employed the AdamW optimizer with binary cross-entropy loss weighted by class frequency, early stopping with a 5-epoch patience window based on validation AUC, and ReduceLROnPlateau learning rate scheduling.

Model performance was assessed using ROC-AUC and average precision (AP) computed from out-of-fold predictions across all five scaffold-aware folds. Per-fold AUC values are reported as mean ± standard deviation across the five folds.

#### Prospective Library Screening and Compound Selection

The Enamine Protein Mimetic Library comprising 8,960 compounds was obtained as a CSV file containing compound identifiers and SMILES strings. All compounds were featurized using Morgan circular fingerprints (radius 2, 2,048 bits) as described above. For each of the four targets, the RF, XGB, and DL models were retrained on the full corresponding training dataset and applied to generate binding probability scores for all 8,960 library compounds. Final compound rankings were generated using an ensemble framework in which predictions from the three models were combined to produce a consensus binding score. Compounds were ranked independently by their target-specific ensemble score, and the top 25 compounds per target were selected for experimental testing, representing a 99.7% reduction in screening burden relative to full-library testing. Compound selection was performed blinded to experimental outcomes.

#### Full Library Random Screening

To establish a rigorous experimental baseline, all 8,960 compounds of the Enamine Protein Mimetic Library were subjected to single-dose (100 μM) binding screening against all four targets using the Dianthus NT.23 Pico system (NanoTemper Technologies) with TRIC technology, following protocols established in our laboratory. Each target protein was fluorescently labeled with RED-tris-NTA 2nd Generation dye at 10 nM and incubated with library compounds at 100 µM in assay buffer (PBS, pH 7.4, 0.05% Tween-20, 2% final DMSO) for 15 minutes at room temperature. Negative controls (assay buffer, n = 8 per replicate) and positive controls (known binders at 100 µM, n = 8 per replicate) were included on each plate. Plates were measured on the Dianthus NT.23 Pico using 5-second laser excitation. Compounds exhibiting ΔF_norm_ values greater than three standard deviations from the negative control mean were designated primary hits. Orthogonal control experiments were conducted for all primary hits to exclude compounds exhibiting autofluorescence, fluorescence quenching, or aggregation artifacts, following established exclusion criteria. Data from two independent experiments were averaged for final hit determination.

#### HTS-Oracle v2 Prospective Screening

The 25 model-selected compounds per target were screened using identical TRIC protocols to the full library random screening baseline, ensuring direct comparability of all experimental measurements. All primary hits from model-selected compounds were subjected to dose-response binding affinity measurements using Monolith X (NanoTemper Technologies) with spectral shift detection. Target proteins were labeled with RED-tris-NTA 2nd Generation dye at 40 nM. Test compounds were prepared from 10 mM DMSO stock solutions and serially diluted. All dose-response measurements were performed in triplicate. Raw thermophoresis data were analyzed using MO.Affinity Analysis software (NanoTemper Technologies), and KD values were determined by nonlinear regression curve fitting using GraphPad Prism 10 with a standard binding isotherm model.

#### Enrichment Factor and Hit Capture Rate Calculation

The enrichment factor for each target was calculated as:

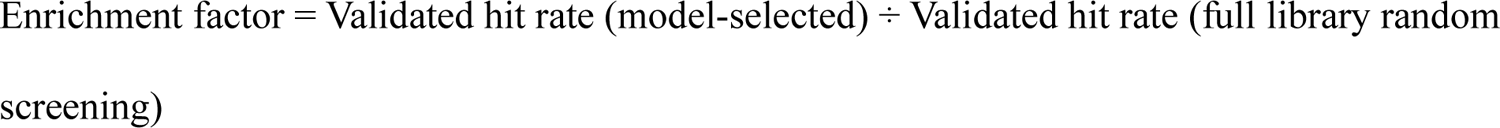

where validated hit rate is defined as the number of dose-response confirmed binders divided by the total number of compounds tested in each group. The hit capture rate was calculated as the number of validated hits identified in the model-selected top 25 compounds divided by the total number of validated hits identified across the entire 8,960-compound library by random screening, expressed as a percentage. Screening burden reduction was calculated as (1 − 25/8,960) × 100%.

#### Functional Blockade Assays

All functional blockade assays were performed using commercially available kits according to the manufacturer’s recommended protocols. CD28 inhibition screening kit was obtained from BPS Bioscience (Catalog# 72007). ICOS inhibition assay was procured from BPS Bioscience (Catalog# 79673). LAG-3 inhibition kit was obtained from Revvity (Catalog# 64LAG3PEG). TIGIT inhibition assay was procured from BPS Bioscience (Catalog# 72029). All experimental runs were performed in triplicates.

#### Statistical Analysis

All reported AUC values represent the mean across five scaffold-aware cross-validation folds, with standard deviation reported as a measure of fold-to-fold variability. ROC curves and precision-recall curves were generated from aggregated out-of-fold predictions across all five folds using scikit-learn. KD values are reported as mean ± standard deviation from triplicate dose-response measurements. GraphPad Prism 10 was used for all dose-response curve fitting and data visualization.

## Data Availability

The code is publicly available at https://github.com/gabr2003/HTS-Oracle-v2.git

