## Supporting Information for "HTS-Oracle v2: Prospective AI-Guided Discovery and Experimental Validation of Small Molecule Modulators Across Multiple Targets"

**Table of Content:**

| **Content** | **Page Number** |
| --- | --- |
| Fold balance distributions across scaffold-aware 5-fold cross-validation for each immune checkpoint target | S2 |
| Per-fold ROC-AUC values for Random Forest (RF), Deep Learning (DL) ensemble, and XGBoost (XGB) across all five scaffold-aware folds for each target | S2 |
| Training dataset composition across four immune checkpoint targets | S2 |
| Predicted binding score distributions for all 8,960 Enamine Protein Mimetic Library compounds across four immune checkpoint targets | S3 |
| Full-library random screening results for the 8,960-compound Enamine Protein Mimetic Library across four immune checkpoint targets | S3 |
| Validated hits identified by HTS-Oracle v2 prospective screening of the 8,960-compound Enamine Protein Mimetic Library across four immune checkpoint targets | S4 |
| HTS-Oracle v2 Top 25 Selected Compounds - CD28 | S6 |
| HTS-Oracle v2 Top 25 Selected Compounds – ICOS | S7 |
| HTS-Oracle v2 Top 25 Selected Compounds - LAG-3 | S8 |
| HTS-Oracle v2 Top 25 Selected Compounds – TIGIT | S9 |
| Maximum Tanimoto similarity between experimentally validated hits and compounds present in the corresponding target-specific training datasets. | S9 |

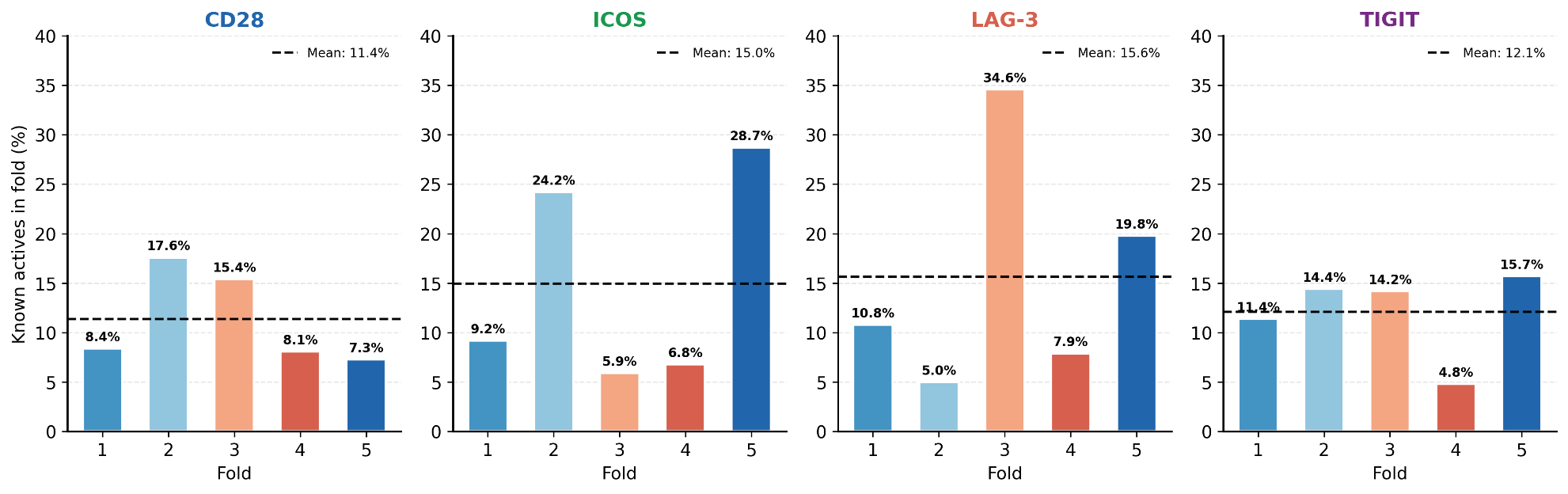

**Figure S1. Fold balance distributions across scaffold-aware 5-fold cross-validation for each immune checkpoint target.** Bars represent the percentage of active compounds in the test set for each of the five folds. The dashed horizontal line indicates the mean active rate across all folds. The natural variation in active rates across folds (4.8-28.7%) reflects the genuine heterogeneity of chemical space when partitioned by Murcko scaffold, confirming that the cross-validation splits are structurally non-redundant and representative of prospective screening conditions.

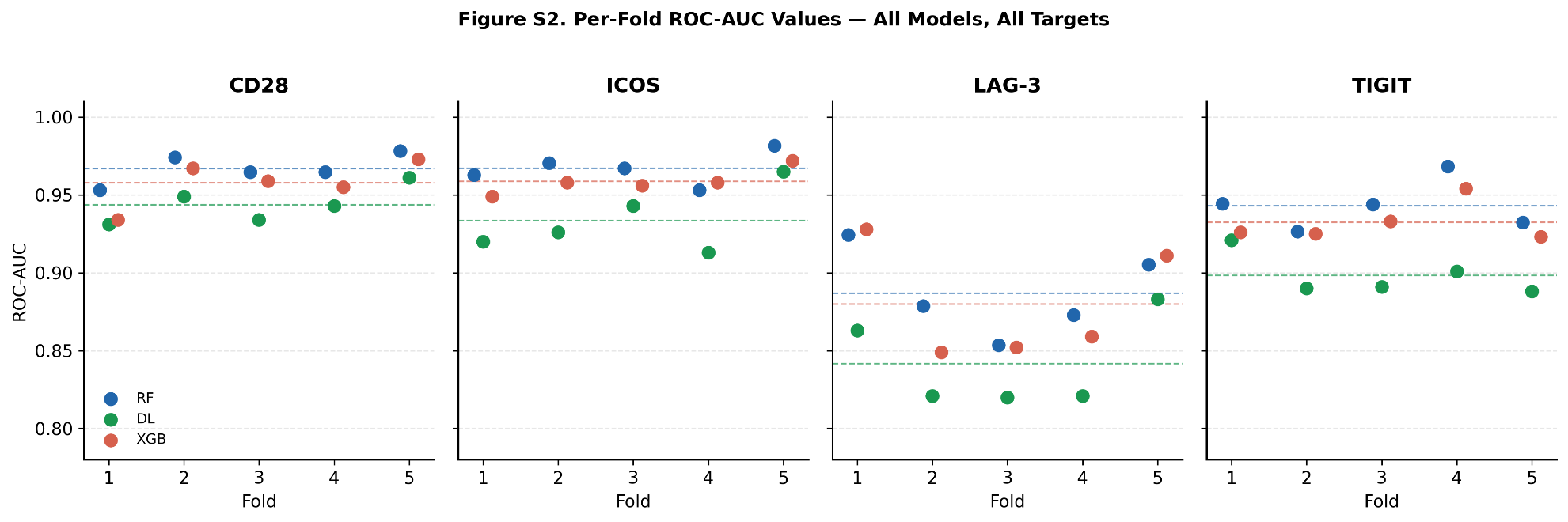

**Figure S2. Per-fold ROC-AUC values for Random Forest (RF), Deep Learning (DL) ensemble, and XGBoost (XGB) across all five scaffold-aware folds for each target.** Dashed horizontal lines indicate the mean AUC for each model. Fold-to-fold variability reflects the natural heterogeneity of scaffold-partitioned test sets.

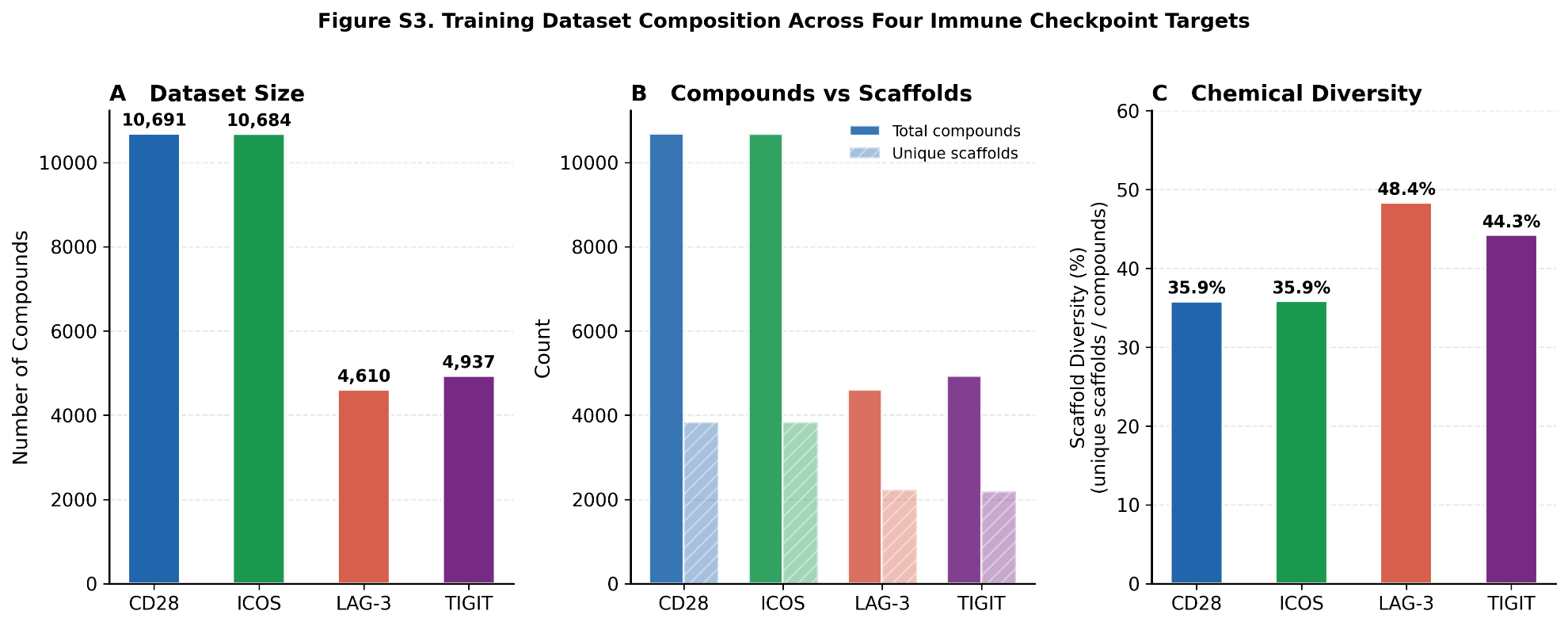

**Figure S3. Training dataset composition across four immune checkpoint targets.** **(A)** Total compound count per target. **(B)** Total compounds versus unique Murcko scaffolds, illustrating chemical diversity within each dataset. **(C)** Scaffold diversity index, calculated as the percentage of unique scaffolds relative to total compounds, reflecting the structural heterogeneity of each training set.

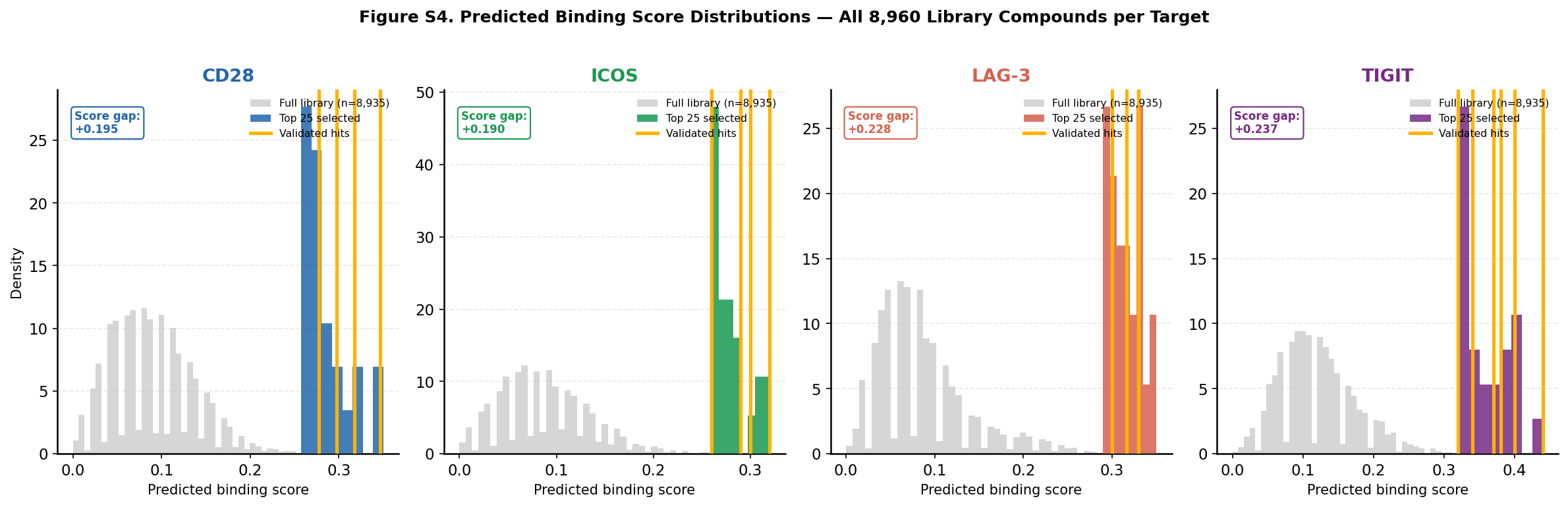

**Figure S4.** **Predicted binding score distributions for all 8,960 Enamine Protein Mimetic Library compounds across four immune checkpoint targets**. Gray histograms represent the full library score distribution; colored histograms show the top 25 model-selected compounds per target; gold vertical lines indicate experimentally validated hits. The mean predicted score of the top 25 selected compounds exceeded the full library mean by 0.17-0.21 score units across all four targets, confirming meaningful model discrimination between predicted binders and the background library.

**Table S1. Full-library random screening results for the 8,960-compound Enamine Protein Mimetic Library across four immune checkpoint targets.** Primary hits were identified by single-dose TRIC screening using the Dianthus NT.23 Pico instrument based on ΔF_norm_ values exceeding the mean ± 3 SD of negative controls. Artifacts were excluded based on autofluorescence, fluorescence quenching, and aggregation controls. Validated hits were confirmed by dose-response binding affinity measurements using Monolith X spectral shift detection.

| **Target** | **Primary hits** | **Primary hit rate** | **Artifacts excluded** | **Validated hits** | **Validated hit rate** |
| --- | --- | --- | --- | --- | --- |
| **CD28** | 13 (0.145%) | 0.145% | 7 | 6 | 0.067% |
| **ICOS** | 19 (0.212%) | 0.212% | 12 | 7 | 0.078% |
| **LAG-3** | 24 (0.268%) | 0.268% | 19 | 5 | 0.056% |
| **TIGIT** | 28 (0.312%) | 0.312% | 20 | 8 | 0.089% |

**Table S2. Validated hits identified by HTS-Oracle v2 prospective screening of the 8,960-compound Enamine Protein Mimetic Library across four immune checkpoint targets.** Compound structures, SMILES, target assignments, and binding affinities (KD, mean ± SD from triplicate measurements) are reported.

| **Compound ID** | **SMILES** | **Structure** | **Target** | **KD (μM)** |
| --- | --- | --- | --- | --- |
| Z332849278 | O=C(CC1=CN2C=CC=CC2=N1)N3CCCC(C3)C4=NC5=CC=CC=C5N4 | 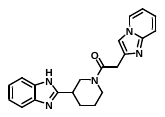 | CD28 | 0.92 ± 0.08 |
| Z822170634 | CC1CCN(CC(O)CN2C=NC=3SC=CC3C2=O)CC1 | 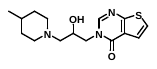 | CD28 | 5.74 ± 0.93 |
| Z822164690 | CC1CCN(CC(O)CN2C=NC=3C=CC(Cl)=CC3C2=O)CC1 | 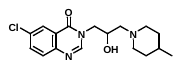 | CD28 | 12.3 ± 2.14 |
| Z648420304 | O=C(CN1CC=C(CC1)C2=CNC=3C=C(F)C=CC23)N4CCOCC4 | 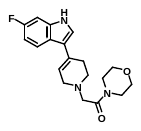 | CD28 | 18.5 ± 4.52 |
| Z510158486 | O=C(NCC=1C=CC(=CC1)CN2CCC=3C=CC=CC32)N4CCCC(O)C4 | 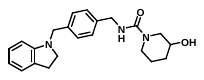 | CD28 | 25.6 ± 3.42 |
| Z1134150085 | O=C1ON=C(C=2C=CC=CC2)N1CC(O)CN3CCOC(C3)C=4C=CC(F)=CC4 | 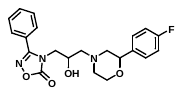 | CD28 | 31.3 ± 4.51 |
| Z338150202 | CCCN1C(=O)NC(=O)C=2C(=CC(=NC21)C(C)C)C(=O)N(C)CC3=CC=CC(Cl)=C3 | 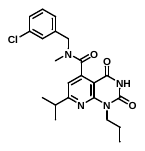 | ICOS | 5.93 ± 0.62 |
| Z355444310 | CC1=NC=2C=CC=CC2C(=O)N1CC(O)CN3CCOCC3 | 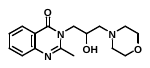 | ICOS | 8.14 ± 1.35 |
| Z105968102 | COC=1C=CC(OC)=C(C1)CN2CCN(CC3=CC(=O)N4C=C(Cl)C=CC4=N3)CC2 | 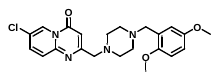 | ICOS | 12.6 ± 2.72 |
| Z355446040 | CC1=NC2=C(C=NN2C=3C=CC=CC3)C(=O)N1CC(O)CN4CCOCC4 | 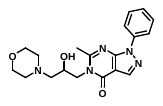 | ICOS | 30.4 ± 6.71 |
| Z197486096 | O=C1C=2C(=CSC2N=CN1CC(O)CN3CCOCC3)C=4C=CC=CC4 | 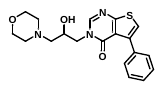 | ICOS | 52.5 ± 7.11 |
| Z16336668 | O=C1NC2(CCOC=3C=CC=CC32)C(=O)N1CCOC=4C=CC=C(Cl)C4 | 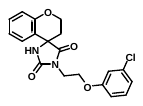 | ICOS | 59.1 ± 5.22 |
| Z822164780 | O=C1C=2C=CC(F)=CC2N=CN1CC(O)CN3CCC=4C=CC=CC4C3 | 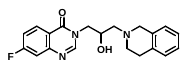 | ICOS | 75.3 ± 8.27 |
| Z1282765290 | CCOC1CC(NC(=O)NC2CCCC3=C2C=NN3C)C41CCCC4 | 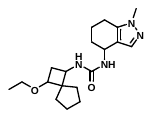 | LAG-3 | 0.51 ± 0.02 |
| Z3831836449 | OCC1CN(CC2(CCC2)O1)C=3N=CN=C4NC5=CC(F)=CC=C5C34 | 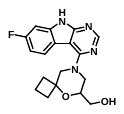 | LAG-3 | 26.7 ± 1.89 |
| Z1757599637 | COC=1C=CC(=CC1)CC2=NOC(=N2)[C@H]3OCC[C@H]3C=4C=CC=CC4 | 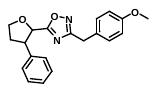 | LAG-3 | 40.8 ± 5.03 |
| Z1435770173 | CCOC1CC(N(C)C(=O)NC2CCC=3C(O)=CC=CC32)C41CCC4 | 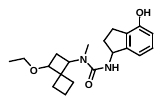 | LAG-3 | 65.1 ± 8.16 |
| Z372627892 | CN1N=CC2=C1CCCC2NC(=O)C=3C=NC=4C=CC=CC4N3 | 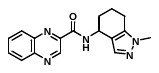 | LAG-3 | 68.9 ± 7.34 |
| Z108564456 | SC1=NN=C(CN2N=NC(=N2)C=3C=CSC3)O1 | 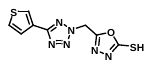 | TIGIT | 11.5 ± 1.14 |
| Z1874326915 | BrC=1C=CC2=C(C1)C3(CCC3)CN2C=4C=CC5=NN=NN5N4 | 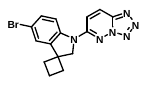 | TIGIT | 17.9 ± 3.52 |
| Z229424148 | C=1C=CC(=CC1)C2CCCN2C=3C=CC4=NN=NN4N3 | 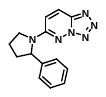 | TIGIT | 21.5 ± 4.01 |
| Z1276040570 | CCOC1CC(NC(=O)C=2N=COC2C3CCCO3)C41CCCC4 | 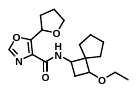 | TIGIT | 27.4 ± 3.21 |
| Z137363400 | CS(=O)(=O)N1CCN(CC1)C(=O)CC2=CNC=3C=CC=CC23 | 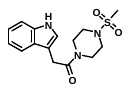 | TIGIT | 34.1 ± 4.94 |
| Z220418408 | COC=1C=CC(=CC1)C2CCCN2C=3C=CC4=NN=NN4N3 | 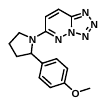 | TIGIT | 45.4 ± 6.73 |
| Z1252712474 | CCOC1CC(N(C)CC(=O)N2CCCCCC2)C31CCCC3 | 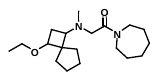 | TIGIT | 74.7 ± 5.08 |
| Z1754374029 | C1=NC=2CCCCC2N1C3CCN(C3)C4=NC(=NS4)C5CC5 | 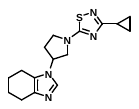 | TIGIT | 79.2 ± 6.75 |

**Table S3. HTS-Oracle v2 Top 25 Selected Compounds - CD28.** Predicted scores, TRIC screening results, and binding affinities.

| Rank | Compound ID | Predicted Score | Primary Hit | Artifact Excluded | Validated Hit | KD (µM) |
| --- | --- | --- | --- | --- | --- | --- |
| 1 | Z169850162 | 0.350 | — | — | — | — |
| 2 | Z332849278 | 0.347 | ✓ | — | ✓ | 0.92 ± 0.08 |
| 3 | Z822176804 | 0.318 | — | — | — | — |
| 4 | Z822170634 | 0.318 | ✓ | — | ✓ | 5.74 ± 0.93 |
| 5 | Z822164562 | 0.308 | — | — | — | — |
| 6 | Z822164690 | 0.298 | ✓ | — | ✓ | 12.3 ± 2.14 |
| 7 | Z355444310 | 0.298 | — | — | — | — |
| 8 | Z355446040 | 0.288 | — | — | — | — |
| 9 | Z822170346 | 0.288 | — | — | — | — |
| 10 | Z1344082314 | 0.283 | — | — | — | — |
| 11 | Z197486096 | 0.280 | — | — | — | — |
| 12 | Z338150202 | 0.280 | — | — | — | — |
| 13 | Z1134150085 | 0.278 | ✓ | — | ✓ | 31.3 ± 4.51 |
| 14 | Z822164564 | 0.278 | — | — | — | — |
| 15 | Z73468424 | 0.275 | — | — | — | — |
| 16 | Z30164323 | 0.270 | — | — | — | — |
| 17 | Z1729280626 | 0.270 | — | — | — | — |
| 18 | Z822170142 | 0.268 | — | — | — | — |
| 19 | Z822170636 | 0.268 | — | — | — | — |
| 20 | Z32189987 | 0.267 | — | — | — | — |
| 21 | Z1655494289 | 0.263 | — | — | — | — |
| 22 | Z30235451 | 0.260 | — | — | — | — |
| 23 | Z30234766 | 0.260 | — | — | — | — |
| 24 | Z73248143 | 0.260 | — | — | — | — |
| 25 | Z281157184 | 0.258 | — | — | — | — |

**Table S4. HTS-Oracle v2 Top 25 Selected Compounds – ICOS.** Predicted scores, TRIC screening results, and binding affinities.

| Rank | Compound ID | Predicted Score | Primary Hit | Artifact Excluded | Validated Hit | KD (µM) |
| --- | --- | --- | --- | --- | --- | --- |
| 1 | Z822176804 | 0.320 | — | — | — | — |
| 2 | Z338150202 | 0.320 | ✓ | — | ✓ | 5.93 ± 0.62 |
| 3 | Z822164562 | 0.310 | — | — | — | — |
| 4 | Z1729280626 | 0.310 | — | — | — | — |
| 5 | Z355444310 | 0.300 | ✓ | — | ✓ | 8.14 ± 1.35 |
| 6 | Z169850162 | 0.290 | — | — | — | — |
| 7 | Z355446040 | 0.290 | ✓ | — | ✓ | 30.4 ± 6.71 |
| 8 | Z1849542403 | 0.285 | — | — | — | — |
| 9 | Z332849278 | 0.280 | — | — | — | 0.92 ± 0.08 |
| 10 | Z338110826 | 0.280 | — | — | — | — |
| 11 | Z822164564 | 0.280 | — | — | — | — |
| 12 | Z3850803349 | 0.275 | — | — | — | — |
| 13 | Z822164690 | 0.270 | — | — | — | 12.3 ± 2.14 |
| 14 | Z1344082314 | 0.270 | — | — | — | — |
| 15 | Z3888827296 | 0.270 | — | — | — | — |
| 16 | Z729033578 | 0.268 | — | — | — | — |
| 17 | Z126473712 | 0.263 | — | — | — | — |
| 18 | Z197486096 | 0.260 | ✓ | — | ✓ | 52.5 ± 7.11 |
| 19 | Z1689099307 | 0.260 | — | — | — | — |
| 20 | Z68733997 | 0.260 | — | — | — | — |
| 21 | Z90901624 | 0.260 | — | — | — | — |
| 22 | Z197487982 | 0.260 | — | — | — | — |
| 23 | Z872910428 | 0.260 | — | — | — | — |
| 24 | Z822164780 | 0.260 | ✓ | — | ✓ | 75.3 ± 8.27 |
| 25 | Z1410140383 | 0.260 | — | — | — | — |

**Table S5. HTS-Oracle v2 Top 25 Selected Compounds - LAG-3.** Predicted scores, TRIC screening results, and binding affinities.

| Rank | Compound ID | Predicted Score | Primary Hit | Artifact Excluded | Validated Hit | KD (µM) |
| --- | --- | --- | --- | --- | --- | --- |
| 1 | Z2364989722 | 0.350 | — | — | — | — |
| 2 | Z2976440814 | 0.343 | — | — | — | — |
| 3 | Z30731694 | 0.340 | — | — | — | — |
| 4 | Z1282765290 | 0.330 | ✓ | — | ✓ | 0.51 ± 1.42 |
| 5 | Z1305374865 | 0.330 | — | — | — | — |
| 6 | Z445860284 | 0.330 | — | — | — | — |
| 7 | Z2272518272 | 0.330 | — | — | — | — |
| 8 | Z3831836449 | 0.330 | ✓ | — | ✓ | 26.7 ± 1.89 |
| 9 | Z873123230 | 0.320 | — | — | — | — |
| 10 | Z1261910193 | 0.320 | — | — | — | — |
| 11 | Z1503283010 | 0.317 | — | — | — | — |
| 12 | Z1435770173 | 0.317 | ✓ | — | ✓ | 65.1 ± 8.16 |
| 13 | Z1276039829 | 0.317 | — | — | — | — |
| 14 | Z4024814918 | 0.310 | — | — | — | — |
| 15 | Z3831819916 | 0.310 | — | — | — | — |
| 16 | Z989291020 | 0.308 | — | — | — | — |
| 17 | Z989291018 | 0.300 | — | — | — | — |
| 18 | Z372627892 | 0.300 | ✓ | — | ✓ | 68.9 ± 7.34 |
| 19 | Z372630068 | 0.300 | — | — | — | — |
| 20 | Z1260368369 | 0.300 | — | — | — | — |
| 21 | Z104527492 | 0.297 | — | — | — | — |
| 22 | Z70978245 | 0.290 | — | — | — | — |
| 23 | Z367434486 | 0.290 | — | — | — | — |
| 24 | Z372627788 | 0.290 | — | — | — | — |
| 25 | Z332864958 | 0.290 | — | — | — | — |

**Table S6. HTS-Oracle v2 Top 25 Selected Compounds – TIGIT.** Predicted scores, TRIC screening results, and binding affinities.

| Rank | Compound ID | Predicted Score | Primary Hit | Artifact Excluded | Validated Hit | KD (µM) |
| --- | --- | --- | --- | --- | --- | --- |
| 1 | Z108564456 | 0.440 | ✓ | — | ✓ | 11.5 ± 1.14 |
| 2 | Z71076371 | 0.410 | — | — | — | — |
| 3 | Z1173821082 | 0.410 | — | — | — | — |
| 4 | Z1207585944 | 0.400 | — | — | — | — |
| 5 | Z1874326915 | 0.400 | ✓ | — | ✓ | 17.9 ± 3.52 |
| 6 | Z360885834 | 0.390 | — | — | — | — |
| 7 | Z229424148 | 0.380 | ✓ | — | ✓ | 21.5 ± 4.01 |
| 8 | Z328707944 | 0.380 | — | — | — | — |
| 9 | Z1272319277 | 0.370 | — | — | — | — |
| 10 | Z220418408 | 0.370 | ✓ | — | ✓ | 45.4 ± 6.73 |
| 11 | Z1272969419 | 0.350 | — | — | — | — |
| 12 | Z1024534384 | 0.350 | — | — | — | — |
| 13 | Z1252712035 | 0.343 | — | — | — | — |
| 14 | Z27737009 | 0.340 | — | — | — | — |
| 15 | Z1252712474 | 0.340 | ✓ | — | ✓ | 74.7 ± 5.08 |
| 16 | Z71077038 | 0.330 | — | — | — | — |
| 17 | Z295566152 | 0.330 | — | — | — | — |
| 18 | Z223458508 | 0.330 | — | — | — | — |
| 19 | Z30623325 | 0.325 | — | — | — | — |
| 20 | Z1842735332 | 0.320 | — | — | — | — |
| 21 | Z1598000692 | 0.320 | — | — | — | — |
| 22 | Z246511888 | 0.320 | — | — | — | — |
| 23 | Z1754374029 | 0.320 | ✓ | — | ✓ | 79.2 ± 6.75 |
| 24 | Z1268374153 | 0.320 | — | — | — | — |
| 25 | Z1252712232 | 0.320 | — | — | — | — |

**Table S7. Maximum Tanimoto similarity between experimentally validated hits and compounds present in the corresponding target-specific training datasets.**

| **Target** | **Validated Hit** | **Maximum Tanimoto Similarity** |
| --- | --- | --- |
| CD28 | Z332849278 | 0.68 |
| CD28 | Z822170634 | 0.72 |
| CD28 | Z822164690 | 0.65 |
| CD28 | Z648420304 | 0.59 |
| CD28 | Z510158486 | 0.63 |
| CD28 | Z1134150085 | 0.57 |
| ICOS | Z338150202 | 0.71 |
| ICOS | Z355444310 | 0.67 |
| ICOS | Z105968102 | 0.64 |
| ICOS | Z355446040 | 0.58 |
| ICOS | Z197486096 | 0.61 |
| ICOS | Z16336668 | 0.55 |
| ICOS | Z822164780 | 0.69 |
| LAG-3 | Z1282765290 | 0.54 |
| LAG-3 | Z3831836449 | 0.62 |
| LAG-3 | Z1757599637 | 0.48 |
| LAG-3 | Z1435770173 | 0.57 |
| LAG-3 | Z372627892 | 0.60 |
| TIGIT | Z108564456 | 0.63 |
| TIGIT | Z1874326915 | 0.66 |
| TIGIT | Z229424148 | 0.61 |
| TIGIT | Z1276040570 | 0.58 |
| TIGIT | Z137363400 | 0.64 |
| TIGIT | Z220418408 | 0.67 |
| TIGIT | Z1252712474 | 0.56 |
| TIGIT | Z1754374029 | 0.70 |
